# Coordination amongst quadriceps muscles suggests neural regulation of internal joint stresses, not simplification of task performance

**DOI:** 10.1101/781534

**Authors:** Cristiano Alessandro, Adarsh Prashara, David P. Tentler, Hsin-Yun Yeh, Filipe O. Barroso, Matthew C. Tresch

## Abstract

Many studies have demonstrated co-variation between muscle activations during behavior, suggesting that muscles are not controlled independently. According to one common proposal, this co-variation reflects simplification of task performance by the nervous system, so that muscles with similar contributions to task variables are controlled together. Alternatively, this co-variation might reflect regulation of low-level aspects of movements that are common across tasks, such as stresses within joints. We examined these issues by analyzing co-variation patterns in quadriceps muscle activity during locomotion in rats. The three mono-articular quadriceps muscles (vastus medialis, VM; vastus lateralis, VL; vastus intermedius, VI) produce knee extension and so have identical contributions to task performance; the bi-articular rectus femoris (RF) produces an additional hip flexion. Consistent with the proposal that muscle co-variation is related to similarity of muscle actions on task variables, we found that the co-variation between VM and VL was stronger than their co-variations with RF. However, co-variation between VM and VL was also stronger than their co-variations with VI. Since all vastii have identical actions on task variables, this finding suggests that co-variation between muscle activity is not solely driven by simplification of task performance. Instead, the preferentially strong co-variation between VM and VL is consistent with the control of internal joint stresses: since VM and VL produce opposing mediolateral forces on the patella, the high positive correlation between their activation minimizes the net mediolateral patellar force. These results provide important insights into the interpretation of muscle co-variations and their role in movement control.

## Introduction

The detailed spatiotemporal structure of muscle activations can provide insight into the control strategies used by the central nervous system (CNS) to produce movement[1–4]. For instance, muscle activations have been shown to co-vary during the production of many behaviors. According to one common hypothesis, the co-variation between muscle activations reflects a neural strategy in which muscles with similar contributions to task performance are controlled together as a single functional unit, often referred to as a ‘muscle synergy’[5–8]. This strategy might simplify task performance by reducing the number of variables that need to be specified for the production of behavior. For example, the quadriceps muscles vastus lateralis (VL) and vastus medialis (VM) have very similar contributions to task performance, with each muscle producing a similar extension torque at the knee[9–11]. Consistent with the proposal that co-variation patterns reflect muscles’ contribution to task performance[7,12–15], the activations of VM and VL are strongly correlated[15–18], suggesting that the CNS might control VM and VL as a single functional unit to simplify the achievement of task goals[19].

Alternatively, co-variation of muscle activations might reflect aspects of motor control other than achieving task goals. Although VM and VL have similar contributions to task performance, they produce opposing mediolateral forces on the patella[11,20,21]. The strong correlation between VM and VL might therefore reflect minimization of net mediolateral patellar forces to prevent aberrant patellofemoral loading[17,22]. In this interpretation, co-variation between these muscles reflects the regulation of low-level biomechanical features that are common across tasks, such as those affecting joint integrity, rather than simplification of task performance.

We evaluated these issues by recording the activity in all four quadriceps muscles, including rectus femoris (RF) and vastus intermedius (VI), during locomotion across a number of task conditions in the rat. Like VM and VL, RF and VI both produce knee extension, but RF also produces an extra flexion torque at the hip (Fig. 1a). If co-variation patterns amongst muscles reflect the similarity of their contributions to task performance, the correlation between the three vastii muscles (VM, VL, VI) should be equally strong and stronger than their correlations to RF since these three muscles all have the same contribution to task performance. However, unlike VM and VL, neither VI nor RF produces a strong mediolateral force on the patella. Therefore, if co-variation patterns amongst muscles reflect regulation of internal joint stresses, the correlation between VM and VL should be higher than the correlation between any other pair of quadriceps muscles, reflecting the need to balance mediolateral forces on the patella. Further, such a low-level control strategy to balance mediolateral patellar forces should be common across variations in task conditions; hence, we expect the correlation between VM and VL to be the highest independent of speed and incline of locomotion. Our results support these latter predictions, suggesting that co-variation patterns amongst quadriceps muscles reflect control of low-level aspects of internal joint mechanics better than simplification of task performance. These results therefore call for a reinterpretation of previous studies examining muscle activation covariation, and suggest the importance of internal joint stresses and strains when investigating neural control strategies.

**Figure 1.**
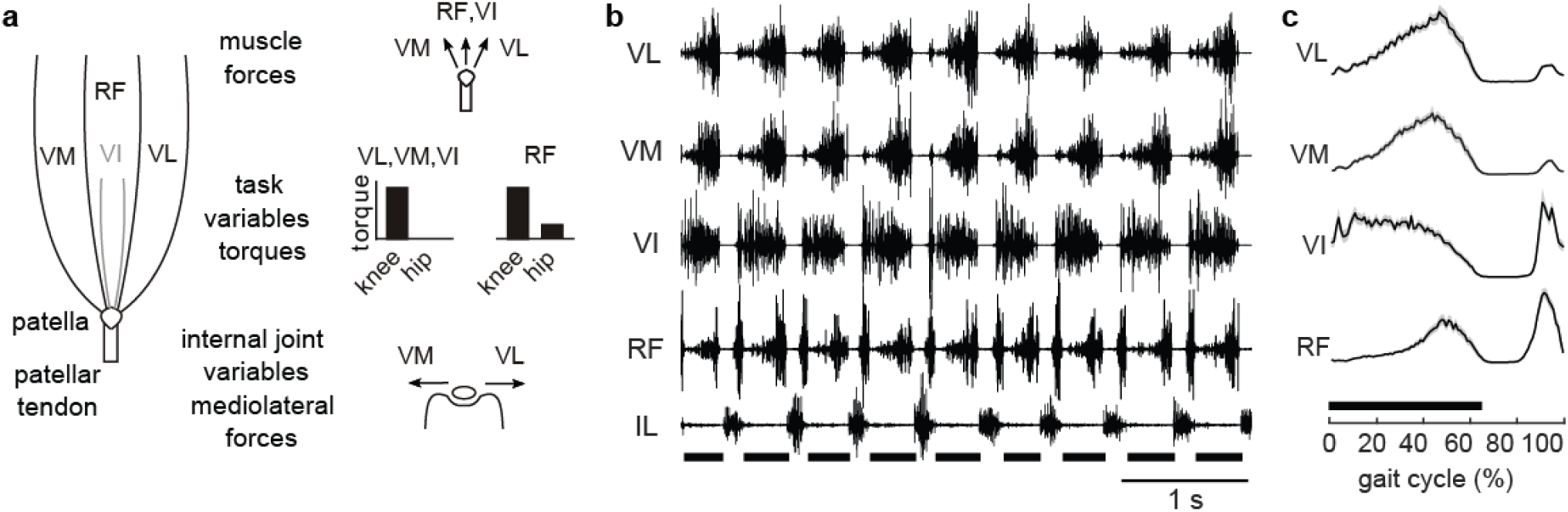
Quadriceps actions and example EMGs recordings. **(a)** Quadriceps muscles attach to the patella, which transfers muscle forces to the tibia via the patellar tendon. Forces from VM, VL, VI and RF produce extension torque at the knee. In addition, RF produces a flexion torque at the hip. Besides joint torques (task variables), VM and VL also produce opposite mediolateral forces on the patella, while RF and VI have minimal effect on mediolateral patellar forces (internal joint variables). Poor regulation of these mediolateral forces might cause aberrant contact stresses within the knee due to patellar loading or displacement. **(b)** Raw EMG signals illustrating the good signal-to-noise ratio obtained in these experiments and the typical patterns of muscle activity for one animal during upslope locomotion at 20 m/min. The black thick horizontal lines at the bottom of the traces indicate the stance phase of locomotion for the implanted limb. Note that the activity of the quadriceps muscles alternates with the activity of the hip flexor illipsoas (IL), which is strongly active during the swing phase of the gait cycle. **(c)** The EMG envelopes of the quadriceps muscles, time-normalized to percentage of gait cycle, illustrate that all the quadriceps muscles are active during the stance phase and in the last part of the swing phase. These activation time courses are represented as mean ± standard error (SE) across strides (number of strides n_s_=99).

## Results

### Stride-averaged activity of VM and VL are strongly correlated across task conditions

We first examined the co-variation patterns amongst stride-averaged activity of each quadriceps muscle across task conditions. If co-variation patterns reflect simplification of task performance, the activity of VM, VL, and VI should be similar to each other for all task conditions but distinct from the activity of RF. If these patterns reflect control of internal joint stresses, the activity in VM and VL should be similar to each other but distinct from activity in both RF and VI.

An example of the activity in quadriceps muscles for one animal and task condition (upslope locomotion at 20 m/min) is illustrated in Fig. 1b, demonstrating the good signal to noise ratio of EMG recordings in these experiments and typical muscle activation patterns. In general, all four quadriceps were active starting in the late portion of the swing phase prior to foot contact and maintained activity throughout the stance phase (Fig. 1c). The activity of VI and RF had slightly different activation profiles, or time courses, from that of VL and VM: VI was highly active during early stance, RF activity was slightly shifted towards late stance, and both VI and RF had a prominent burst of activity during swing.

This general pattern of muscle activity was similar across animals and task conditions (Fig. 2a). For all speeds and inclines of locomotion, the average activity of quadriceps muscles across animals started in late swing and was maintained throughout the stance phase. However, the detailed time course (i.e. the activation profile or envelope of activation) and intensity (i.e. the overall level of activation) of each muscle activation were different across conditions. In some behavioral conditions, the activity of multiple quadriceps muscles was very similar; e.g. VM, VL, and RF each had similar activity patterns during downslope walking. Strikingly, only the activity patterns in VM and VL remained similar across all task conditions.

**Figure 2.**
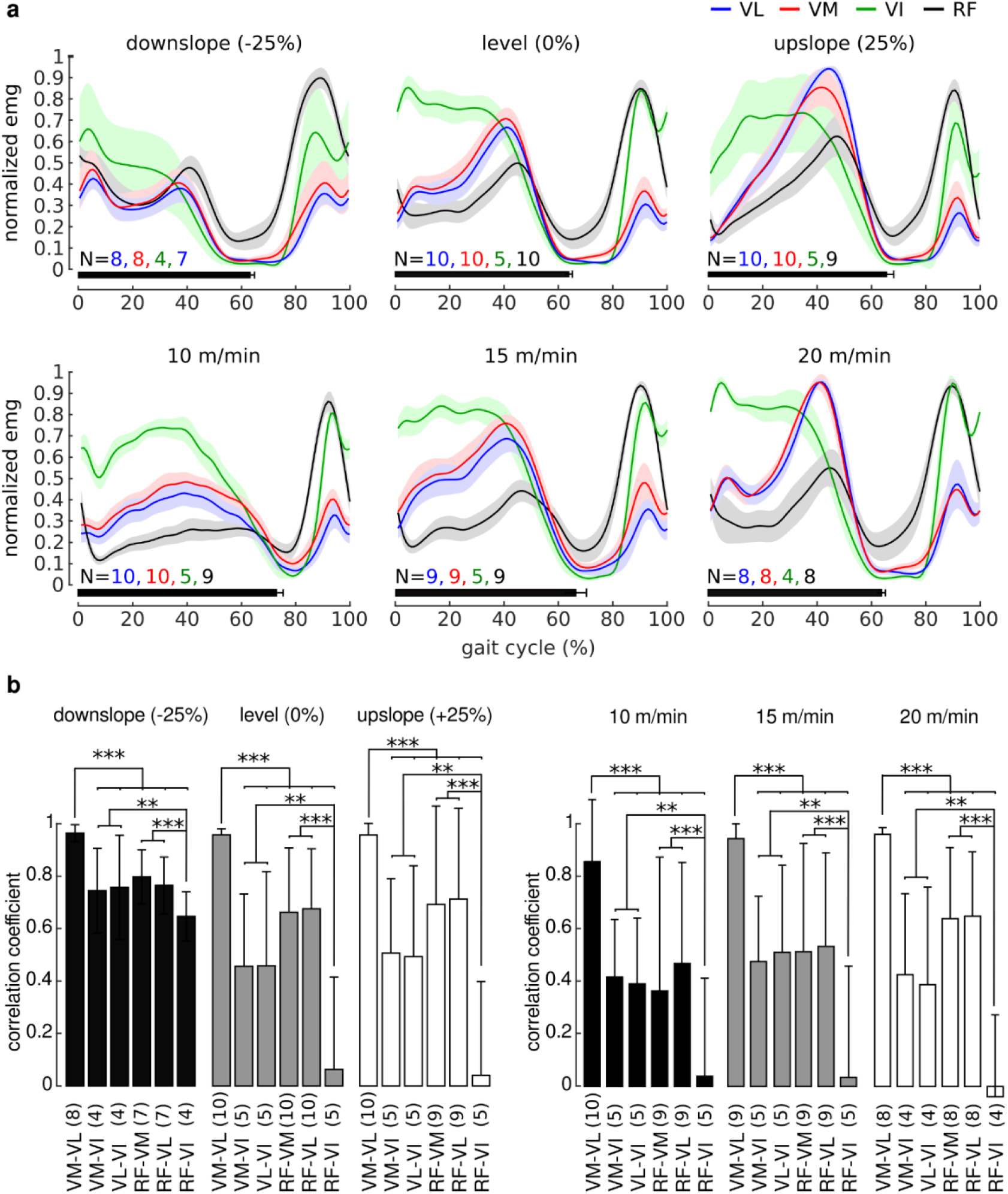
Stride-averaged activity of VM and VL are strongly coordinated across task conditions. **(a)** Stride-averaged quadriceps muscle activity across all animals is modulated by speed and incline of locomotion, both in terms of activation time course and in terms of activation intensity. However, the averaged activities of VM and VL are strikingly similar to each other for all task conditions, suggesting a tight coordination between these two muscles. Signals are represented as mean±SE across animals (number of animals as indicated in the image). Signals from each animal are averaged across strides (n_s_=113±60, mean ± standard deviation, SD, across animals and conditions). Note that the number of animals varies across conditions and muscles due to the inclusion criteria described in the Methods section. The activation of each muscle is normalized by its maximum across speeds or inclines for each animal for display purposes; all analyses described in the Results were performed on the unnormalized values. **(b)** Correlation between stride-averaged activation time courses of VM and VL is stronger than correlation between stride-averaged activation time courses of any other pair of muscles for each behavioral condition. Data are shown as mean±SD across animals (number of animals as indicated in the bars; average number of strides for each animal as indicated above). Significance levels: * * p<0.01; * * * p<0.001.

Statistical analyses of stride-averaged activation time courses confirmed these observations (Fig. 2b), demonstrating strong correlation between VM and VL in all behavioral conditions (r_incline_=0.96±0.03, p<0.001; r_speed_=0.92±0.15, p<0.001). Most importantly, these correlations were higher than the correlations between the stride-averaged activation time courses of any other pair of muscles including those involving VI (all comparisons, p<0.001) for all behavioral conditions (ANOVA, p_incline:muscle_=0.61, p_speed:muscle_=0.75). We also found that the correlations between stride-averaged activation time courses of RF and VI were the lowest for all inclines (p_VMVL=VLVI_=0.003; p_VMVL=VMVI_=0.006; other comparisons: p<0.001) and speeds (p_VMVL=VLVI_=0.002; p_VMVL=VMVI_=0.002; other comparisons: p<0.001), reflecting the observation that VI was more active during early stance whereas RF was more active during late stance.

### Correlation between VM and VL time courses on individual strides is strongest for all task conditions

If the CNS coordinates quadriceps muscles in order to balance mediolateral forces on the patella, not only should the activation time courses of VM and VL be similar on average (Fig. 2) but the time courses of these two muscles should be similar on each individual stride. We therefore evaluated the correlations between the activation time courses of each pair of muscles for each stride of locomotion. Figure 3 illustrates the distributions of the correlation coefficients, for one animal and locomotor condition, calculated for each individual stride. In this example, the activation time courses of VM and VL were highly correlated for the vast majority of the strides. Further, VM and VL were more strongly correlated than all other muscle pairs. Note also that the correlation between RF and VI appeared to be lower than the correlation between other muscle pairs, although there was high variability across strides.

**Figure 3.**
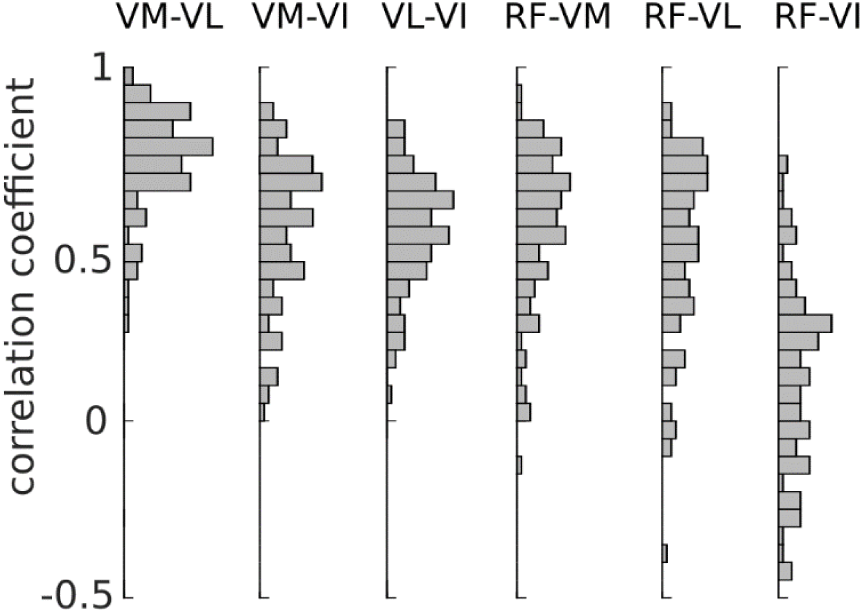
Correlations between activation time courses of quadriceps muscle activity on individual strides for a representative animal in one task condition. The distributions of the correlation coefficients (n_s_=101) between the activation time courses of quadriceps muscles for each stride demonstrate a very tight coordination between VM and VL for this animal and behavioral condition. The correlation between VM and VL is higher that the correlations between the time courses of any other pair of muscles. The correlation between RF and VI is the weakest on average, with a large variability that also includes negative correlation coefficients.

These co-variation patterns were consistent across animals and task conditions (Fig. 4). The time courses of VM and VL activation were highly correlated on individual strides for all locomotor conditions (r_speed_=0.82±0.07, p<0.001; r_incline_=0.86±0.04, p<0.001), and significantly more correlated than those of any other muscle pair for all inclines (p_VMVL=VLVI,−25%_=0.02; p_VMVL=VMVI,−25%_=0.006; other comparisons: p<0.001) and for all speeds (all comparisons: p<0.001). These results demonstrate that the correlation between VM and VL was consistently the strongest correlation amongst quadriceps muscles independent of task conditions, supporting the idea that co-variation patterns reflect regulation of internal joint stresses rather than simplification of task performance.

**Figure 4.**
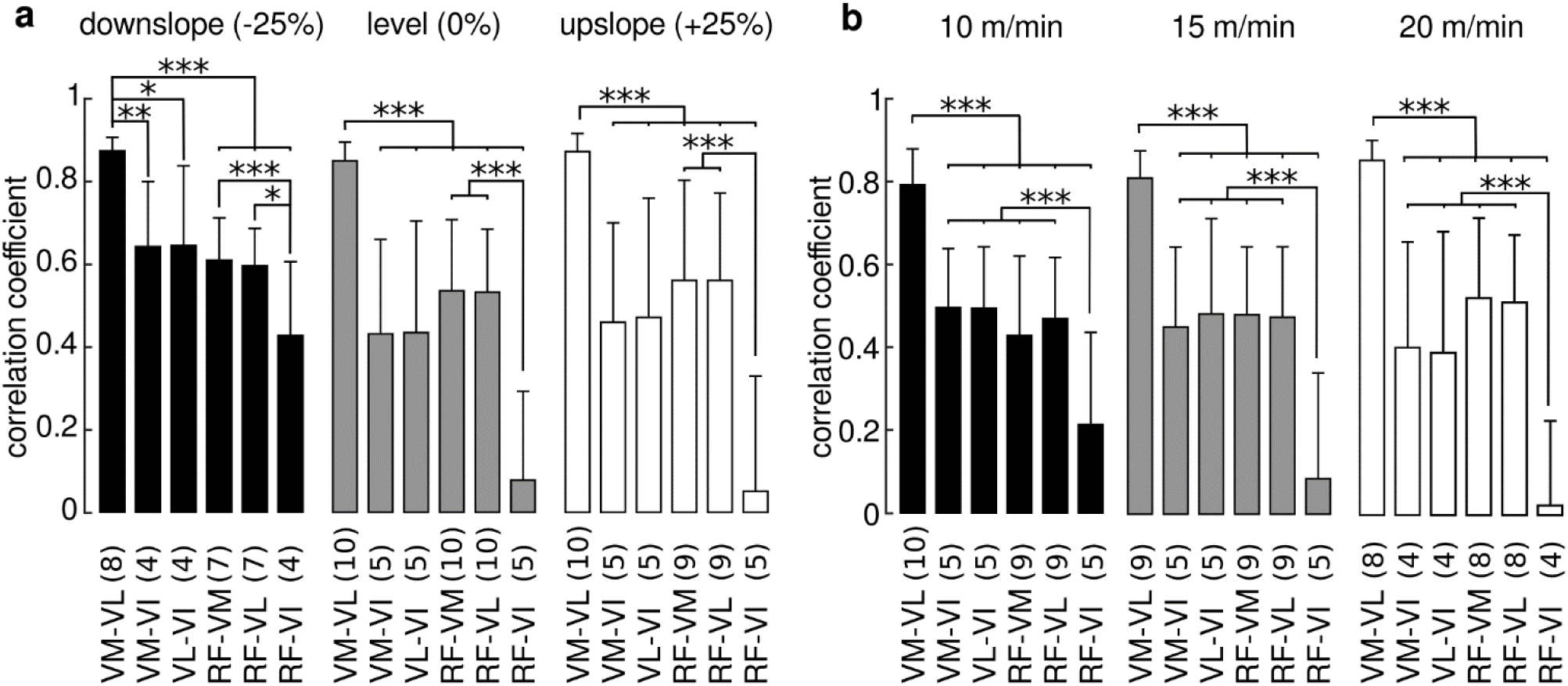
Correlation between VM and VL time courses on individual strides is strongest for all task conditions. The correlation between the activation time courses of VM and VL on individual strides is significantly higher than the correlation between the time courses of any other pair of muscles. This result is independent of incline (**a**) and speed (**b**) of locomotion. Correlation between RF and VI is the lowest for all speeds, and lower than RF-VL and RF-VM for all inclines. Data are represented as mean ± SD across animals (number of animals as indicated in the bars; number of strides for each animal, incline: 121±66, speed: 105±56). Significance levels: * p<0.05; * * p<0.01; * * * p<0.001.

Similar to the results analyzing stride-averaged muscle activity (Fig. 2), we also found that the correlation between the activation time courses of RF and VI on individual strides was lower than the correlations between RF and VM (p_-25%_<0.001; p_0%_<0.001; p_25%_<0.001) and RF and VL (p_-25%_=0.01; p_0%_<0.001; p_25%_<0.001) for all inclines, and lower than the correlations between any other muscle pair for all speeds (all comparisons: p<0.001).

### Correlation between VL and VM activation intensities across strides is strongest for all task conditions

The analysis of muscle activation time course described in the previous section does not account for co-variation in activation intensities across strides: two muscles can have similar activation time courses in each individual stride, but different patterns of intensity co-variation across strides (Fig. 5a). If the CNS coordinates the activity of VM and VL, the intensity in these muscles should co-vary. We therefore analyzed the correlation between activation intensities of each pair of quadriceps muscles across strides for each task condition. Figure 5b illustrates the results of this analysis for an animal in one task condition, showing that the correlation between activation intensities of VM and VL across strides was stronger than the correlation between the intensities of any other pair of muscles.

**Figure 5.**
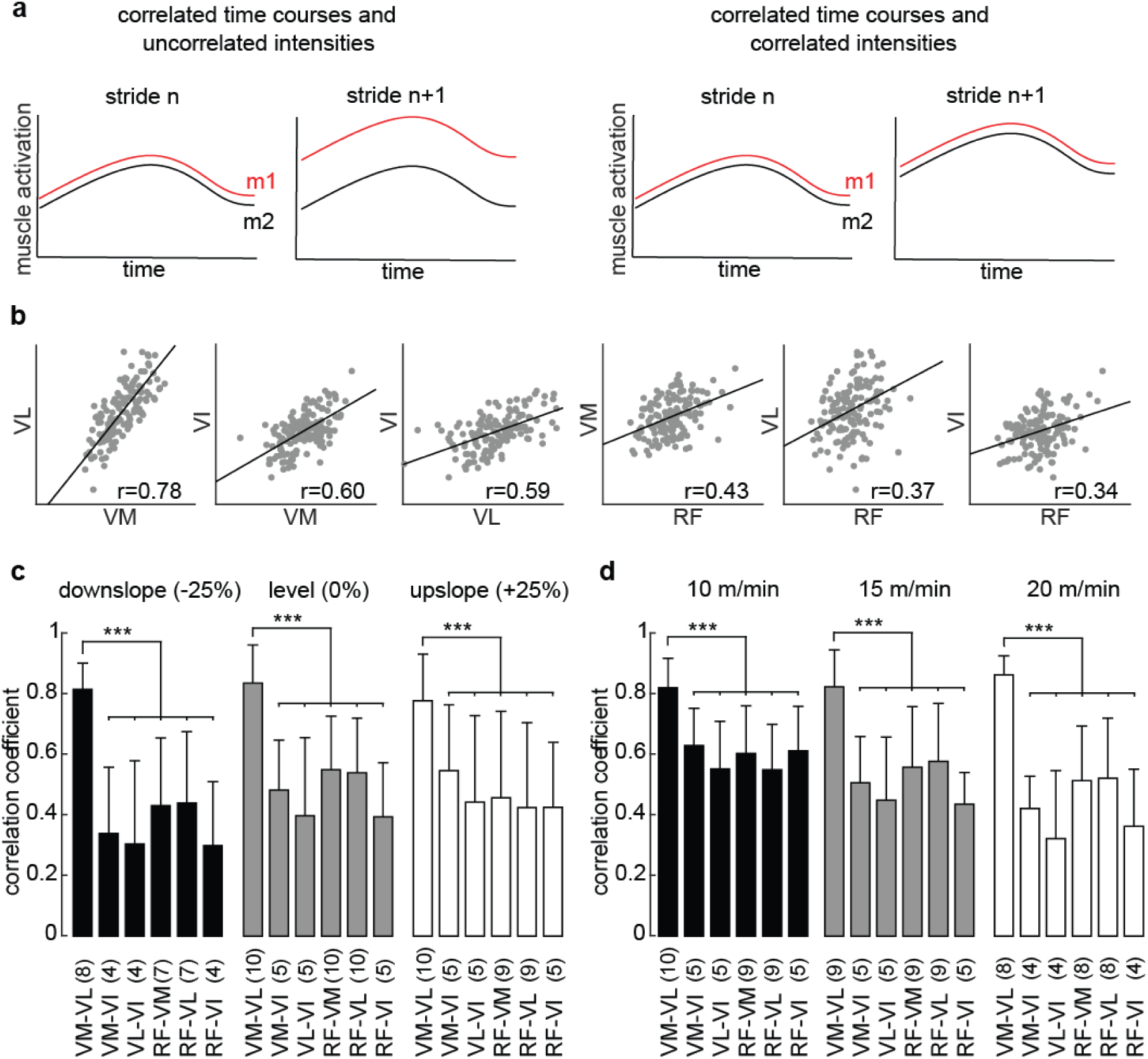
Correlation between activation intensity across strides. **(a)** High correlation between activation time courses within the gait cycle does not imply high correlation between activation intensity across strides. The cartoon on the left illustrates an example in which the activation intensities of two muscles (m1 and m2) change independently from each other from one stride to the next, although their activation time courses are highly correlated within the gait cycle. The cartoon on the right shows muscles that have both correlated activation intensities and correlated activation time courses. Panel **(b)** illustrates the activation intensities of each pair of quadriceps muscles for each stride of locomotion (each represented by a dot, n_s_=141) and corresponding correlation coefficients (r), for a representative animal and behavioral condition. In this example, the activation intensities of VM and VL are strongly correlated, and more strongly correlated than the activation intensities of the other muscle pairs. **(c**,**d)** The correlation between the activation intensities of VM and VL across strides is significantly higher than the correlation between any other muscle pair. This result holds for all inclines (**c**) and speeds (**d**) of locomotion. The correlations between the other pairs of muscles were not significantly different from one another. Data are represented as mean ± standard deviations (SD) across animals (number of animals as indicated in the bars; number of strides for each animal, incline: 121±66, speed: 105±56). * * * p<0.001.

Similar results were obtained across animals and behavioral conditions (Fig. 5c). The correlation between the activation intensities of VM and VL was strong (r_incline_=0.81±0.13, p<0.001; r_speed_=0.83±0.10, p<0.001) and was higher than the correlations between all other muscle pairs (all comparisons: p<0.001). The higher correlation between VM and VL was consistent across behavioral conditions (ANOVA, p_incline:muscle_=0.97, p_incline_=0.37, p_speed:muscle_=0.30, p_speed_=0.14). The correlations amongst the activation intensities of the other muscle pairs were not significantly different from one another for all inclines (all comparison: p=1) and speeds (p_VMRF=VLVI_=0.74; all other comparisons: p=1). These results again demonstrate the preferentially strong co-variation between VM and VL activation across task conditions.

## Discussion

We examined the patterns of co-variation amongst quadriceps muscles during locomotion in the rat. The correlation between the activity of VM and VL was stronger than the correlation between the activity of any other pair of quadriceps muscles, and this stronger correlation was consistent across behavioral conditions. In particular, the activities of VM and VL were more strongly correlated than the activity of either of these muscles and VI, even though these three muscles have the same action on task performance. This higher correlation between VM and VL was observed for the stride-averaged time courses of muscle activity, for the time courses of muscle activity on each individual strides, and for the co-variation of muscle intensities across strides. These results demonstrate that co-variation patterns amongst quadriceps muscles do not simply reflect simplification of task performance; rather, these results support the idea that these co-variations reflect regulation of internal joint mechanics.

### Evaluating muscle co-variation patterns

We evaluated muscle co-variation in terms of the similarity of activation time courses and overall intensities of muscle activity in this study. We used these measures because of their functional interpretability, evaluating whether sets of muscles are activated in similar ways across a variety of task conditions. Although the relationship between EMG and muscle force is complex[23], these measures can also provide information about mechanical consequences of muscle activation strategies; i.e. about the contributions of these muscles to task performance (their torques across joints) and their contributions to internal joint mechanics (their mediolateral forces on the patella)

Other measures of muscle co-variation have been used to provide insight into the strategies underlying the neural control of movement. Dimensionality reduction analyses have been used to identify muscle co-variation patterns (‘muscle synergies’) amongst large numbers of muscles in order to evaluate coordination strategies across entire limbs and bodies[5,8,24–26]. The measures of muscle co-variation we used in this study are clearly related to those analyses, since both reflect co-variation of EMG time courses and intensities and, in fact, quadriceps muscles often appear as a single synergy when analyzing EMGs during locomotion[27,28]. Although dimensionality reduction techniques provide insights into neural control strategies, they can be difficult to perform (e.g. choosing the correct number of synergies) and to interpret (e.g. whether important information is left in the residual variance). The measures we used in this study are more straightforward to perform and to interpret in terms of motor function.

A different approach for evaluating muscle co-variation is to examine the precise timing of motor units in different muscles, using either time domain cross-correlations or frequency domain coherence analyses[1,15,16]. These measures provide important information about the neural drive to motor units, evaluating whether they share common inputs from the nervous system. Of particular relevance to the current study, previous experiments showed that there is strong coherence in the activity of motor units in VM and VL [15]. Although this result is consistent with the strong correlations in activity time courses and intensities observed here, it is important to note that the two measures are not necessarily related: it is possible to have correlated activations without having strong coherence, and to have strong coherence without correlated activity. Although there are difficulties in recording motor unit activity in free behaviors such as locomotion especially for deep muscles such as VI, it would be interesting to evaluate coherence between motor units in VI and the other quadriceps muscles to determine whether the correlation patterns observed here are mirrored at the level of individual motor units.

Finally, we note that although the measures of muscle coordination we considered here are related to each other, they reflect different aspects of muscle activation patterns and provide complementary information. For example, in many behavioral conditions the activation time courses of RF and VI appeared temporally shifted relative to one another (Fig. 2), with VI more strongly activated during early stance and RF more strongly activated during late stance. These distinct activity patterns resulted in correlations between the activation time courses of RF and VI that were lower than those between other muscle pairs (Figs. 2, 4). However, the correlation between the intensities of RF and VI was similar to the correlation between any other muscle pair (other than VM-VL, Fig. 5), suggesting that although these muscles have distinct time courses, they have similar modulation of intensity across strides. The fact that for all measures of muscle co-variation considered here, the correlation between VM and VL was stronger than that between any other muscles demonstrates the robustness of this result.

### Functional role of muscle co-variation in quadriceps

Our results provide important insights about the functional interpretation of the co-variation between quadriceps muscle activity. Although previous studies have demonstrated a strong correlation between the activity of VM and VL[10,29–31], the functional role of this correlation has not been clear. Since VM and VL have the same contribution to task variables and produce opposing mediolateral patellar forces[11], their strong correlation is consistent with both simplification of task performance[16,19] and regulation of internal joint stresses[29,32,33]. Similarly, the observation that RF could be controlled more independently than the vasti[31,34] is also consistent with both interpretations, as RF has a different task action from VM and VL[35,36] and produces minimal mediolateral patellar force (Fig. 1a). In order to distinguish between these competing interpretations, we recorded the activity in VI. This muscle has the same task action as VM and VL, but has minimal action on mediolateral patellar force[37]. Our observation that the correlation between VM and VL was higher than the correlations of either muscle to VI provides strong support that co-variation patterns between muscles best reflect regulation of internal joint stresses. It is important to note that although the correlation between VM and VL was consistently high in these experiments, it was not perfect. Whether this residual variation in VM and VL activation has functionally relevant consequences or is simply intrinsic variability is unclear.

Our results are consistent with previous work suggesting that the CNS actively coordinates quadriceps activity so as to minimize the net mediolateral force on the patella[17,22,38–42]. Although EMG recordings provide an indirect estimate of muscle force[23], the strong correlations between the time course and intensity of VM and VL observed here would be expected to limit the net mediolateral patellar force. Consistent with this idea, alterations in the coordination between VM and VL have been associated with the development of patellofemoral pain or patellar maltracking[17,43]. These observations suggest that co-variation patterns between muscles, and their alteration, might provide insights into the development of musculoskeletal disorders and suggest interventions strategies to improve rehabilitation outcomes.

A more independent control of VI from the other quadriceps would allow the CNS to regulate other aspects of motor control while balancing VL and VM to minimize patellar loading. For example, VI could be used to stabilize the knee joint during flexion by providing antagonistic co-contractions to the hamstrings [44], consistent with the high level of VI activity we observed during the swing phase of locomotion in all behavioral conditions (Fig. 2a). Similarly, VI might be used to compensate for reductions in RF activity during fatiguing contractions[18]. This strategy is consistent with the idea of uncontrolled manifolds[2], in which the activity of RF and VI would be negative correlated in order to obtain a stable knee torque, although we did not observe such negative correlations between RF and VI here. Alternatively, activation of VI might help patellar stabilization but in a manner distinct from the co-activation of VM and VL: the relatively constant activation of VI throughout the stance phase of locomotion might ensure the patella is locked firmly within the patellar groove, thereby preventing patellar dislocation. Additional experiments will be necessary to clarify the functional role of VI and its relationship to other quadriceps muscles.

The observation that the strong VM-VL correlation was consistent across task conditions might suggest that these patterns are produced by lower-level sensorimotor systems, potentially within the spinal cord. For example, last order spinal interneurons might branch to activate both VM and VL motoneurons[15,45] or be strongly coordinated by feedback from sensory afferents[46] conveying information about patellar loading via joint receptors[47,48] or muscle proprioceptors[49,50]. However, previous work has suggested that the strong co-variation between VM and VL motor units is due to both cortical and spinal systems[15,16], suggesting that this co-variation may result from distributed processing in multiple areas of the CNS. Further work will be needed to understand the neural systems involved in coordinating these muscles and ensuring joint integrity.

Note that a low-level, spinal implementation of VM-VL coordination for regulation of internal joint mechanics might indirectly simplify task performance. By ensuring proper control of joint integrity, it would reduce the number of criteria that have to be specified by higher-level systems involved in performing tasks and achieving behavioral goals. This reduction of complexity would be similar to the potential simplification of muscle mechanics by spinal reflexes[49] and is consistent with previous suggestions of hierarchical control strategies in the CNS[51].

### Interpretation of muscle co-variation patterns in other systems

It will be interesting to evaluate whether the results of these experiments extend to other joints and limb structures. Our ability to disambiguate between interpretations of muscle co-variation relied on the conceptual simplicity of the knee biomechanics in the rat. The rat knee joint provides a clear separation between the action of quadriceps muscles on task performance and internal joint stresses, allowing us to distinguish between the different interpretations of muscle co-variation[11,22]. In other joints and animals such a separation may not be as clear, and the actions of muscles on these different aspects of motor control may need to be estimated using detailed biomechanical models[52,53].

It is also possible that co-variation patterns amongst muscles might reflect different processes depending on the joint/limb being considered. For instance, co-variation patterns amongst finger muscles in primates, identified using motor unit coherence analyses, have been suggested to mainly reflect direct cortical inputs to motor units and appear to vary with task conditions[54]. It will be interesting in future work to reconsider co-variation patterns at other joints and limbs, evaluating whether they reflect regulation of task performance or internal joint mechanics. Regardless of the generalization of these results across limbs and joints, the results of this study, in combination with previous work[55,56], highlights the potential importance of internal joint mechanics when interpreting muscle co-variation patterns and neural control strategies.

## Acknowledgments

This research was supported by the NIH grant number NS086973 (MCT).

## Author Contributions

CA, FOB and MCT conceptualized this study. CA, AP and DT performed the experiments. CA and MCT performed surgeries. CA analyzed the data. CA, AP, DT and HYY processed the data. CA and MCT wrote the manuscript; all the authors revised the manuscript. MCT provided supervision and secured funding.

## Declaration of Interests

The authors declare no competing interests.

## Methods

We performed experiments on adult female Sprague Dawley rats (n=10, weight=0.31±0.02 g). All procedures were approved by the Animal Care Committee of Northwestern University.

### Experimental protocol

We first trained rats to maintain stable gait during treadmill locomotion. We then implanted chronic EMG electrodes in hindlimb muscles and allowed animals to recover for at least 10 days. We recorded EMG activity during treadmill locomotion at different speeds (10, 15 and 20 m/min during level locomotion) and inclines (−25, level, and 25%, at 20 m/min), for a total of 5 behavioral conditions for each rat. The order of these behavioral conditions was randomized across rats in order to avoid biases. At the end of data collection, animals were euthanized and electrode location verified (no misplaced electrodes were found). We also measured the electrical impedance of each electrode, and we excluded from the analysis electrodes with impedance higher than 50 kohm (potentially indicating a damaged wire) and those with values close to zero (indicating a short circuit). Because of these criteria, we only excluded the electrode associated to VI in one animal.

### Implantation of EMG electrodes

The procedure to implant the EMG electrodes was described in detail previously[22,57]. Briefly, we anesthetized animals with isoflurane (3% in O2 ∼2 l/min), shaved their hindlimb, and prepared them for aseptic surgery. We implanted pairs of electrodes in the quadriceps muscles: vastus lateralis (VL), vastus medialis (VM), rectus femoris (RF), and vastus intermedius (VI). Other muscles (up to 8 additional) were implanted in the same hindlimb but were not analyzed in this study. Knots placed on both sides of the muscle secured the exposed electrode sites within the muscle belly. The electrode leads were tunneled subcutaneously to a connector (Omnetics nano series) on the back of the animal. We implanted the deep muscle VI by separating the anterior head of biceps femoris from VL to expose the femur and gently lifting VL from the bone; VI was then clearly distinguishable from VL due to its distinct color and fiber organization. During the first two days after surgery, we administered analgesics (buprenorphine, 0.2 mg/kg, twice daily; meloxicam, 0.25 mg, once daily).

### EMG recordings

Before each recording session, we briefly anesthetized the animal under isoflurane (2-3% in O2 ∼2 l/min) in order to attach the EMG connector in the rat to the amplifier via cable. We also attached retroreflective markers to the skin of the animal to measure hindlimb kinematics, as described previously[22]. We then placed the animal on the treadmill, and waited at least 30 minutes before starting data collection. We recorded at least two minutes of locomotion for each of the behavioral conditions described above (see *Experimental protocol*), and allowed the animal to rest for at least two minutes after each bout of locomotion. Two animals refused to walk downslope (−25%) and were only able to maintain speed of 15 m/min during upslope locomotion (+25% incline). To assess the effect of incline in those animals, we therefore used the data recorded at (15 m/min, 0%) and (15 m/min, 25%) in order to match speed across the different incline conditions.

### Data acquisition and processing

Differential EMG signals were amplified (1000X, A-M Systems Inc., model 3500), band-pass (30-1000Hz) and notch filtered (60Hz), and then digitized (5000Hz, Vicon Lock+, model VL0143). The digitized signals were further high-pass filtered offline to remove motion artifacts (50Hz, 4th order Butterworth).

To assess potential cross-talk among EMG signals, we computed the cross-correlation between the unrectified activity of adjacent muscles. We discarded signals for which the absolute value of the peak cross-correlation was higher than 0.3 [58,59]. This only happened for one rat between muscles VM and VI. Note also that we found distinct activation time courses across behavioral conditions for adjacent quadriceps muscles (see Fig. 2a), further confirming minimal contributions of cross-talk to these recordings. Further, the pair of muscles with the strongest correlation (VM-VL, see Results) was not adjacent to one another. We rectified the remaining signals, and computed their envelopes by low pass filtering (20Hz, 4th order Butterworth).

We segmented the EMG envelopes into separate strides, defining the beginning of each stride as the moment of foot-strike (i.e. when the foot touched the ground, as determined from the trajectory of the toe marker). To obtain consistent data for steady locomotion in each behavioral condition, we only considered strides with durations within 1.5 standard deviations from the mean duration; this criterion eliminated strides in which the animal either accelerated or decelerated across the treadmill. We also excluded strides with clear EMG artifacts that could occur when the cable hit the side of the treadmill, as identified using Tukey outlier analysis (i.e. identifying EMG values that were 1.5 interquartiles above the upper quartile of the maxima across steps). Application of these inclusion criteria resulted in data sets with an average of 110 strides (minimum of 30) for each behavioral condition.

### Measures of muscle coordination

We considered two measures of the coordination between quadriceps muscle activity. The first measure evaluated the similarity between the activation time courses of muscles within the gait cycle. To calculate this activation time course we time-normalized each step so that it consisted of 100 samples. We then computed the Pearson correlation between the activation profiles of all six possible pairs of quadriceps muscles (VM-VL, VM-VI, VL-VI, RF-VM, RF-VL, RF-VI). This measure quantifies how similarly the activity of each pair muscles is modulated across the locomotor cycle. We calculated this correlation two ways: using the stride-averaged activation of each muscle, and using the activation of each muscle on individual strides. The second measure evaluated the co-variation of the overall activation intensities of muscles across strides. To calculate this activation intensity we integrated the time course of each muscle on each stride during the stance phase (i.e. from foot-strike to foot-off, when each quadriceps muscles is consistently activated). We then calculated the Pearson correlation between the activation intensities across strides of each pair of muscles for each behavioral condition.

### Statistical analyses

We employed Linear Mixed Effect Models (LMEM) to perform all the statistical analyses, using the glmmTMB package[60] in the R environment[61]. LMEMs allowed us to exploit the large number of strides for each animal and behavioral condition, obtaining the maximum statistical power from the dataset. Furthermore, LMEMs allowed us to analyze samples of different size (e.g. unequal numbers of strides across behavioral conditions and animals), to cope with missing data, and to take into account variability at different levels of the dataset (e.g. across strides, behavioral conditions, and subjects). After fitting the LMEMs, we tested our specific hypotheses of interest by performing post-hoc tests on the model parameters. We performed planned comparisons with the null hypothesis that the correlation between VM and VL (for both measures) was equal to the correlations between each of the other pairs of muscles for each behavioral condition. Furthermore, we performed multiple post-hoc comparisons to test the difference between the correlations between all other muscle pairs. In both cases, we used two-tail z-tests and we adjusted the p-values using Bonferroni corrections. Due to the high number of tests (all comparisons across muscle pairs), this correction could result in adjusted p-values equal to 1. We considered tests to be statistically significant if their p-values were lower than the 0.05 significance level. Prior to fitting the LMEMs to the data, we transformed the Pearson correlation coefficients with the Fisher z-transform (i.e. inverse hyperbolic tangent function). This transformation renders the sample distribution of Pearson correlation coefficients approximately normal, and therefore allows us to compute confidence intervals and to perform statistical comparisons[62,63]. To confirm that our dataset met the assumption of Gaussian distribution and independence of residuals and random effects[64], we visually inspected the distributions using qq-plots and histograms.

We evaluated the effect of speed and the effect of incline on the measure of similarity amongst activation time courses in two separate LMEM analyses. In both analyses, we used the correlation coefficients as the dependent variable, and the factors *muscle-pair* and *behavioral condition* (either the different levels of speed, or the different levels of incline), as well as their interaction term, as independent variables (i.e. fixed effects). Furthermore, we considered *animal* as random effect. Since there are multiple observations (strides) for each combination of the two factors, we should theoretically employ a “maximal model” with random intercept, slopes and interaction term[65]. In practice, these models are difficult to fit due to the large amount of data required to reliably estimate all the random parameters. Therefore, we followed recent recommendations to simplify these maximal models by: (1) removing potentially irrelevant random effects; (2) constraining the variance-covariance matrix between the random effects; and (3) removing non-significant interaction terms between fixed-effects. This process results in the “parsimonious model” that best fit the data[66]. Starting with a model with random interaction term (which is strictly required to specify that there are multiple observations for each combination of the two factors[67]), we iteratively added random intercept and slopes until the fitting performance (evaluated using Akaike information criterion, AIC) did not improve significantly. Using this iterative procedure, we obtained a model with uncorrelated random intercept, random slope on *muscle-pair* (with diagonal var-cov matrix), and random slope on the interaction between *muscle-pair* and *behavioral condition* (with compound symmetry var-cov matrix), both for speed and incline. We then evaluated the significance of the fixed effects by performing an ANOVA with marginal sum-of-squares on the fitted model; the interaction term was significant for both model and therefore we did not remove it. We performed post-hoc tests to evaluate the difference between the correlation coefficients between pairs of muscles within the same behavioral condition.

For the analysis of similarity between stride-averaged activation time courses of muscle activity, we first calculated the average activation time course of each muscle across strides for each animal and behavioral condition. We then calculated the Pearson correlation coefficients between each pair of these average time courses, for each animal and behavioral condition. Finally, we fit two LMEMs to the Fischer transformed correlation coefficients: one to evaluate the effects of speed, and another to evaluate the effect of incline. We used *muscle-pair* and *behavioral condition* (either the different levels of speed, or the different levels of incline), as well as their interaction term, as fixed effects, and *animal* as random effect. Since this dataset contains a single observation for each combination of *muscle-pair* and *behavioral condition* (i.e. the correlation between stride-averaged activation time courses), the model can only have random intercept, and the var-cov matrix becomes a single variance[64]. We evaluated the significance of the fixed effects by performing an ANOVA with marginal sum-of-squares on the fitted model, and we removed potentially non-significant interaction terms. We performed post-hoc tests as explained above.

Similarly, we fit two LMEMs to the Fischer transformed Pearson correlation coefficients of the activation intensities. We used *muscle-pair* and *behavioral condition*, as well as their interaction term, as fixed effects, and *animal* as random effect. We evaluated the significance of the fixed effects by performing an ANOVA, and we performed post-hoc tests as explained above.

## Data and code availability

The data and code generated during this study are available upon request from the corresponding author.

## References

1. Farina, D., Merletti, R., and Enoka, R.M. (2014). The extraction of neural strategies from the surface EMG: An update. J. Appl. Physiol. 117, 1215–1230.

2. Latash, M., Scholz, J., and Schoener, G. (2007). Toward a new theory of motor synergies. Motor Control 11, 276–308.

3. Valero-Cuevas, F.J. (2015). Fundamentals of Neuromechanics (Springer-Verlag London).

4. Alessandro, C., Backers, N., Goebel, P., Resquin, F., Gonzalez, J., and Osu, R. (2016). Motor Control and Learning Theories. In Emerging Therapies in Neurorehabilitation II, J. Pons, R. Raya, and J. Gonzalez, eds. (Springer International Publishing AG Switzerland), pp. 225–250.

5. Tresch, M., and Jarc, A. (2009). The case for and against muscle synergies. Curr. Opin. Neurobiol. 19, 601–607.

6. Alessandro, C., Ioannis, D., Nori, F., Panzeri, S., and Berret, B. (2013). Muscle synergies in neuroscience and robotics: from input to task-space perspectives. Front. Comput. Neurosci. 7.

7. Ting, L.H., Chiel, H.J., Trumbower, R.D., Allen, J.L., McKay, J.L., Hackney, M.E., and Kesar, T.M. (2015). Neuromechanical principles underlying movement modularity and their implications for rehabilitation. Neuron 86, 38–54.

8. Giszter, S.F. (2015). Motor primitives-new data and future questions. Curr. Opin. Neurobiol. 33, 156–165.

9. De Ruiter, C.J., Hoddenbach, J.G., Huurnink, A., and De Haan, A. (2008). Relative torque contribution of vastus medialis muscle at different knee angles. Acta Physiol. 194, 223–237.

10. Visscher, R.M.S., Rossi, D., Friesenbichler, B., Dohm-Acker, M., Rosenheck, T., and Maffiuletti, N.A. (2017). Vastus medialis and lateralis activity during voluntary and stimulated contractions. Muscle and Nerve 56, 968–974.

11. Sandercock, T.G., Wei, Q., Dhaher, Y.Y., Pai, D.K., and Tresch, M.C. (2018). Vastus lateralis and vastus medialis produce distinct mediolateral forces on the patella but similar forces on the tibia in the rat. J. Biomech. 81, 45–51.

12. Alessandro, C., and Nori, F. (2012). Identification of Synergies by Optimization of Trajectory Tracking Tasks. In Fourth IEEE RAS/EMBS International Conference on Biomedical Robotics and Biomechatronics (Rome: IEEE), pp. 924–930.

13. Alessandro, C., Carbajal, J.P., and D’Avella, A. (2014). A computational analysis of motor synergies by dynamic response decomposition. Front. Comput. Neurosci. 7.

14. Delis, I., Hilt, P.M., Pozzo, T., Panzeri, S., and Berret, B. (2018). Deciphering the functional role of spatial and temporal muscle synergies in whole-body movements. Sci. Rep. 8.

15. Laine, C.M., Martinez-Valdes, E., Falla, D., Mayer, F., and Farina, D. (2015). Motor neuron pools of synergistic thigh muscles share most of their synaptic input. J. Neurosci. 35, 12207–12216.

16. De Marchis, C., Severini, G., Castronovo, A.M., Schmid, M., and Conforto, S. (2015). Intermuscular coherence contributions in synergistic muscles during pedaling. Exp. Brain Res. 233, 1907–1919.

17. Pal, S., Besier, T.F., Draper, C.E., Fredericson, M., Gold, G.E., Beaupre, G.S., and Delp, S.L. (2012). Patellar tilt correlates with vastus lateralis:vastus medialis activation ratio in maltracking patellofemoral pain patients. J. Orthop. Res. 30, 927–933.

18. Akima, H., Saito, A., Watanabe, K., and Kouzaki, M. (2012). Alternate muscle activity patterns among synergists of the quadriceps femoris including the vastus intermedius during low-level sustained contraction in men. Muscle and Nerve 46, 86–95.

19. Mohr, M., Nann, M., Von Tscharner, V., Eskofier, B., and Nigg, B.M. (2015). Task-dependent intermuscular motor unit synchronization between medial and lateral Vastii muscles during dynamic and isometric squats. PLoS One 10.

20. Wilson, N.A., and Sheehan, F.T. (2010). Dynamic in vivo quadriceps lines-of-action. J. Biomech. 43, 2106–2113.

21. Lin, F., Wang, G., Koh, J.L., Hendrix, R.W., and Zhang, L.-Q. (2004). In vivo and noninvasive three-dimensional patellar tracking induced by individual heads of quadriceps. Med. Sci. Sports Exerc. 36, 93–101.

22. Alessandro, C., Rellinger, B.A., Barroso, F.O., and Tresch, M.C. (2018). Adaptation after vastus lateralis denervation in rats demonstrates neural regulation of joint stresses and strains. Elife 7, e38215.

23. Hug, F., Hodges, P.W., and Tucker, K. (2015). Muscle force cannot be directly inferred from muscle activation: Illustrated by the proposed imbalance of force between the vastus medialis and vastus lateralis in people with patellofemoral pain. J. Orthop. Sports Phys. Ther. 45, 360–365.

24. d’Avella, A., Saltied, P., and Bizzi, E. (2003). Combinations of muscle synergies in the construction of a natural motor behavior. Nat. Neurosci. 6, 300–308.

25. Alessandro, C., Delis, I., Nori, F., Panzeri, S., and Berret, B. (2013). Muscle synergies in neuroscience and robotics: from input-space to task-space perspectives. Front. Comput. Neurosci. 7.

26. Torricelli, D., Barroso, F., Coscia, M., Alessandro, C., Lunardini, F., Bravo Esteban, E., and d’Avella, A. (2016). Muscle synergies in clinical practice: Theoretical and practical implications. In Biosystems and Biorobotics, pp. 251–272.

27. Ivanenko, Y.P., Cappellini, G., Dominici, N., Poppele, R.E., and Lacquaniti, F. (2005). Coordination of locomotion with voluntary movements in humans. J. Neurosci. 25, 7238–53.

28. Ivanenko, Y.P., Poppele, R.E., and Lacquaniti, F. (2004). Five basic muscle activation patterns account for muscle activity during human locomotion. J. Physiol. 556, 267–82.

29. Mellor, R., and Hodges, P. (2005). Motor unit synchronization between medial and lateral vasti muscles. Clin. Neurophysiol. 116, 1585–1595.

30. Brøchner Nielsen, N.P., Tucker, K., Dorel, S., Guével, A., and Hug, F. (2017). Motor adaptations to local muscle pain during a bilateral cyclic task. Exp. Brain Res. 235, 607–614.

31. Hug, F., Hodges, P.W., Hoorn, W. Van Den, and Tucker, K. (2014). Between-muscle differences in the adaptation to experimental pain. 1132–1140.

32. Ng, G.Y.F., Zhang, A.Q., and Li, C.K. (2008). Biofeedback exercise improved the EMG activity ratio of the medial and lateral vasti muscles in subjects with patellofemoral pain syndrome. J. Electromyogr. Kinesiol. 18, 128–133.

33. Mellor, R., and Hodges, P.W. (2005). Motor unit syncronization is reduced in anterior knee pain. J. Pain 6, 550–558.

34. Place, N., Matkowski, B., Martin, A., and Lepers, R. (2006). Synergists activation pattern of the quadriceps muscle differs when performing sustained isometric contractions with different EMG biofeedback. Exp. Brain Res. 174, 595–603.

35. Greene, E. (1963). Anatomy of the rat (American Philosophical Society).

36. Johnson, W.L., Jindrich, D.L., Roy, R.R., and Reggie Edgerton, V. (2008). A three-dimensional model of the rat hindlimb: Musculoskeletal geometry and muscle moment arms. J. Biomech. 41, 610–619.

37. Blazevich, A.J., Gill, N.D., and Zhou, S. (2006). Intra- and intermuscular variation in human quadriceps femoris architecture assessed in vivo. J. Anat. 209, 289–310.

38. Fagan, M.K., and Pisoni, D.B. (2009). Perspectives on multisensory experience and cognitive development in infants with cochlear implants. Scand. J. Psychol. 50, 457–462.

39. Keet, J.H.L., Gray, J., Harley, Y., and Lambert, M.I. (2007). The effect of medial patellar taping on pain, strength and neuromuscular recruitment in subjects with and without patellofemoral pain. Physiotherapy 93, 45–52.

40. Ryan, C.G., and Rowe, P.J. (2006). An electromyographical study to investigate the effects of patellar taping on the vastus medialis/vastus lateralis ratio in asymptomatic participants. Physiother. Theory Pract. 22, 309–315.

41. Christou, E.A. (2004). Patellar taping increases vastus medialis oblique activity in the presence of patellofemoral pain. J. Electromyogr. Kinesiol. 14, 495–504.

42. Ng, G.Y.F., and Cheng, J.M.F. (2002). The effects of patellar taping on pain and neuromuscular performance in subjects with patellofemoral pain syndrome. Clin. Rehabil. 16, 821–827.

43. Fagan, V., and Delahunt, E. (2008). Patellofemoral pain syndrome: a review on the associated neuromuscular deficits and current treatment options. Br. J. Sports Med. 42, 789–795.

44. Saito, A., Watanabe, K., and Akima, H. (2013). The highest antagonistic coactivation of the vastus intermedius muscle among quadriceps femoris muscles during isometric knee flexion. J. Electromyogr. Kinesiol. 23, 831–837.

45. Farina, D., and Negro, F. (2015). Common synaptic input to motor neurons, motor unit synchronization, and force control. Exerc. Sport Sci. Rev. 43, 23–33.

46. Jankowska, E. (1992). Interneuronal relay in spinal pathways from proprioceptors. Prog. Neurobiol. 38, 335–378.

47. Iles, J.F., Stokes, M., and Young, A. (1990). Reflex actions of knee joint afferents during contraction of the human quadriceps. Clin. Physiol. Funct. Imaging 10, 489–500.

48. Sjölander, P., Johansson, H., and Djupsjöbacka, M. (2002). Spinal and supraspinal effects of activity in ligament afferents. J. Electromyogr. Kinesiol. 12, 167–176.

49. Wilmink, R.J.H., and Nichols, T.R. (2003). Distribution of heterogenic reflexes among the quadriceps and triceps surae muscles of the cat hind limb. J. Neurophysiol. 90, 2310–24.

50. Blecher, R., Heinemann-Yerushalmi, L., Assaraf, E., Konstantin, N., Chapman, J.R., Cope, T.C., Bewick, G.S., Banks, R.W., and Zelzer, E. (2018). New functions for the proprioceptive system in skeletal biology. Philos. Trans. R. Soc. B Biol. Sci. 373.

51. Loeb, G.E., Brown, I.E., and Cheng, E.J. (1999). A hierarchical foundation for models of sensorimotor control. Exp. Brain Res. 126, 1–18.

52. Lenhart, R.L., Kaiser, J., Smith, C.R., and Thelen, D.G. (2015). Prediction and Validation of Load-Dependent Behavior of the Tibiofemoral and Patellofemoral Joints During Movement. Ann. Biomed. Eng. 43, 2675–2685.

53. Smith, C.R., Won Choi, K., Negrut, D., and Thelen, D.G. (2018). Efficient computation of cartilage contact pressures within dynamic simulations of movement. Comput. Methods Biomech. Biomed. Eng. Imaging Vis. 6, 491–498.

54. Laine, C.M., and Valero-Cuevas, F.J. (2017). Intermuscular coherence reflects functional coordination. J. Neurophysiol. 118, 1775–1783.

55. Solomonow, M., and Krogsgaard, M. (2001). Sensorimotor control of knee stability. A review. Scand. J. Med. Sci. Sports 11, 64–80.

56. O’connor, B.L., Visco, D.M., Brandt, K.D., Albrecht, M., and O’connor, A.B. (1993). Sensory nerves only temporarily protect the unstable canine knee joint from osteoarthritis. evidence that sensory nerves reprogram the central nervous system after cruciate ligament transection. Arthritis Rheum. 36, 1154–1163.

57. Tysseling, V.M., Janes, L., Imhoff, R., Quinlan, K.A., Lookabaugh, B., Ramalingam, S., Heckman, C.J., and Tresch, M.C. (2013). Design and evaluation of a chronic EMG multichannel detection system for long-term recordings of hindlimb muscles in behaving mice. J. Electromyogr. Kinesiol. 23, 531–539.

58. Loeb, G.E., and Gans, C. (1986). Electromyography for Experimentalists.

59. Overduin, S.A., d’Avella, A., Roh, J., and Bizzi, E. (2008). Modulation of muscle synergy recruitment in primate grasping. J. 28, 880–892.

60. Magnusson, A., Skaug, H.J., Nielsen, A., Berg, C., Kristensen, K., Maechler, M., van Bentham, K., Bolker, B.M., and Brooks, M.E. (2017). glmmTMB: generalized linear mixed models using template model builder. R Packag. version 0.1 0.2.3.

61. R Development Core Team (2017). R Development Core Team. R A Lang. Environ. Stat. Comput.

62. Bond, C.F., and Richardson, K. (2004). Seeing the Fisher Z-transformation. In Psychometrika, pp. 291–303.

63. Fischer, R. (1921). On the “probable error” of a coefficient of correlation deduced from a small sample. Metron 1, 3–32.

64. Pinheiro, J.C., and Bates, D.M. (2000). Mixed effects models in S and S-Plus (Springer).

65. Barr, D.J., Levy, R., Scheepers, C., and Tily, H.J. (2013). Random effects structure for confirmatory hypothesis testing: Keep it maximal. J. Mem. Lang. 68, 255–278.

66. Matuschek, H., Kliegl, R., Vasishth, S., Baayen, H., and Bates, D. (2017). Balancing Type I error and power in linear mixed models. J. Mem. Lang. 94, 305–315.

67. Barr, D.J. (2013). Random effects structure for testing interactions in linear mixed-effects models. Front. Psychol. 4, 3–4.

